# fastVEP: A High-Performance Variant Effect Predictor Implemented in Rust

**DOI:** 10.64898/2026.04.14.718452

**Authors:** Kuan-lin Huang

**Author notes:** Corresponding author. Website: https://PrecisionOmics.org.

## Abstract

Annotating genomic variants with their predicted functional consequences is a required step in genomics research and clinical diagnostics. The Ensembl Variant Effect Predictor (VEP) is the community standard for this task, but its Perl implementation struggles with the variant call sets that modern sequencing studies produce. Here we present fastVEP, a complete reimplementation of the VEP variant annotation engine in Rust, a systems programming language that combines memory safety with C-level performance.

fastVEP annotates the complete GIAB HG002 clinical whole-genome sequencing benchmark (4.05 million high-confidence variants) against the full Ensembl GRCh38 gene model (508,530 transcripts) in 86 seconds. Across five organisms and complete gold-standard call sets, including 26 million Mouse Genomes Project variants and 12.9 million Arabidopsis 1001 Genomes variants, throughput holds between 47,000 and 86,000 variants per second. In single-threaded head-to-head runs on the same whole genome, fastVEP is 22-24x faster than Ensembl VEP v115.1, which needs approximately 77 minutes for the equivalent consequence-only run. Accuracy was checked field by field against Ensembl VEP release 115.1: all 23 compared annotation fields matched on 2,340 shared transcript-allele pairs.

Beyond core consequence prediction, fastVEP provides a supplementary annotation framework (fastSA) with a native binary format for direct integration with ClinVar, gnomAD, dbSNP, COSMIC, 1000 Genomes, TOPMed, and MitoMap. It also supplies prediction and conservation scores including PhyloP, GERP, REVEL, SpliceAI, PrimateAI, and SIFT/PolyPhen via dbNSFP; structural variant annotation (DEL, DUP, INV, CNV, BND) with SV-specific consequence prediction; gene-level annotation from OMIM and gnomAD gene constraint metrics; a filter_vep-compatible expression filter engine; multi-sample genotype parsing; regulatory region detection; and mitochondrial-specific variant handling.

fastVEP supports both GRCh38 and GRCh37, predicts 49 Sequence Ontology consequence terms, writes 47 VEP-compatible CSQ annotation fields, generates HGVS nomenclature, and outputs VCF, tab-delimited, or JSON. It ships as a single 3.3 MB statically linked binary with no external dependencies and includes a built-in web interface for interactive annotation; a hosted server is available at https://fastVEP.org

fastVEP is open source under the Apache 2.0 license at https://github.com/Huang-lab/fastVEP

## 1. Introduction

Identifying and interpreting genomic variants is central to modern genomics research and precision medicine. As sequencing costs have fallen, individual whole-genome and whole-exome studies now routinely produce call sets of millions of variants (The 1000 Genomes Project Consortium, 2015; Karczewski et al., 2020). Before downstream analysis and clinical interpretation can begin, every variant needs a predicted functional consequence: does it change an amino acid, disrupt a splice site, fall in a regulatory region, or do nothing at all?

The Ensembl Variant Effect Predictor (VEP) has become the de facto standard for this step (McLaren et al., 2016). VEP predicts the effects of variants on genes, transcripts, and protein sequences using Sequence Ontology (SO) terms (Eilbeck et al., 2005). It reads and writes multiple formats, generates HGVS nomenclature (den Dunnen et al., 2016), and can be extended through plugins.

VEP’s Perl implementation, however, carries costs that matter at scale. The runtime typically needs 500 MB of memory or more, module loading makes startup slow, and single-threaded offline throughput is limited to hundreds of variants per second. These costs are harder to absorb as call sets reach millions of variants in population-scale studies (Taliun et al., 2021) and as clinical laboratories press for faster turnaround.

Several tools address performance directly, including SnpEff (Cingolani et al., 2012), ANNOVAR (Wang et al., 2010), Nirvana (Stromberg et al., 2024), and echtvar (Pedersen, 2023). echtvar in particular showed that Rust’s systems-level performance, combined with compact integer encoding and binary search, can reach million-variant-per-second annotation for population frequency lookups. Each tool makes its own trade-offs in annotation completeness, SO term coverage, and compatibility with the VEP ecosystem.

Here we present fastVEP, a complete reimplementation of the VEP consequence prediction engine in Rust (Matsakis & Klock, 2014). Rust guarantees memory safety without garbage collection, giving C/C++ performance with the safety of managed languages. fastVEP sustains 47,000 to 86,000 variants per second across five model organisms and annotates a complete human whole genome 22-24x faster than Ensembl VEP v115.1 in single-threaded head-to-head runs. It does so while matching VEP on all 23 validated annotation fields, shipping as a dependency-free static binary, and including a built-in web interface for interactive use.

## 2. Design and Implementation

### 2.1 Architecture Overview

fastVEP is organized as a Cargo workspace of twelve modular crates (libraries), following Rust’s model of composable, independently testable components (**Figure 1**). The implementation totals approximately 43,000 lines of Rust source code.

**Figure 1.**
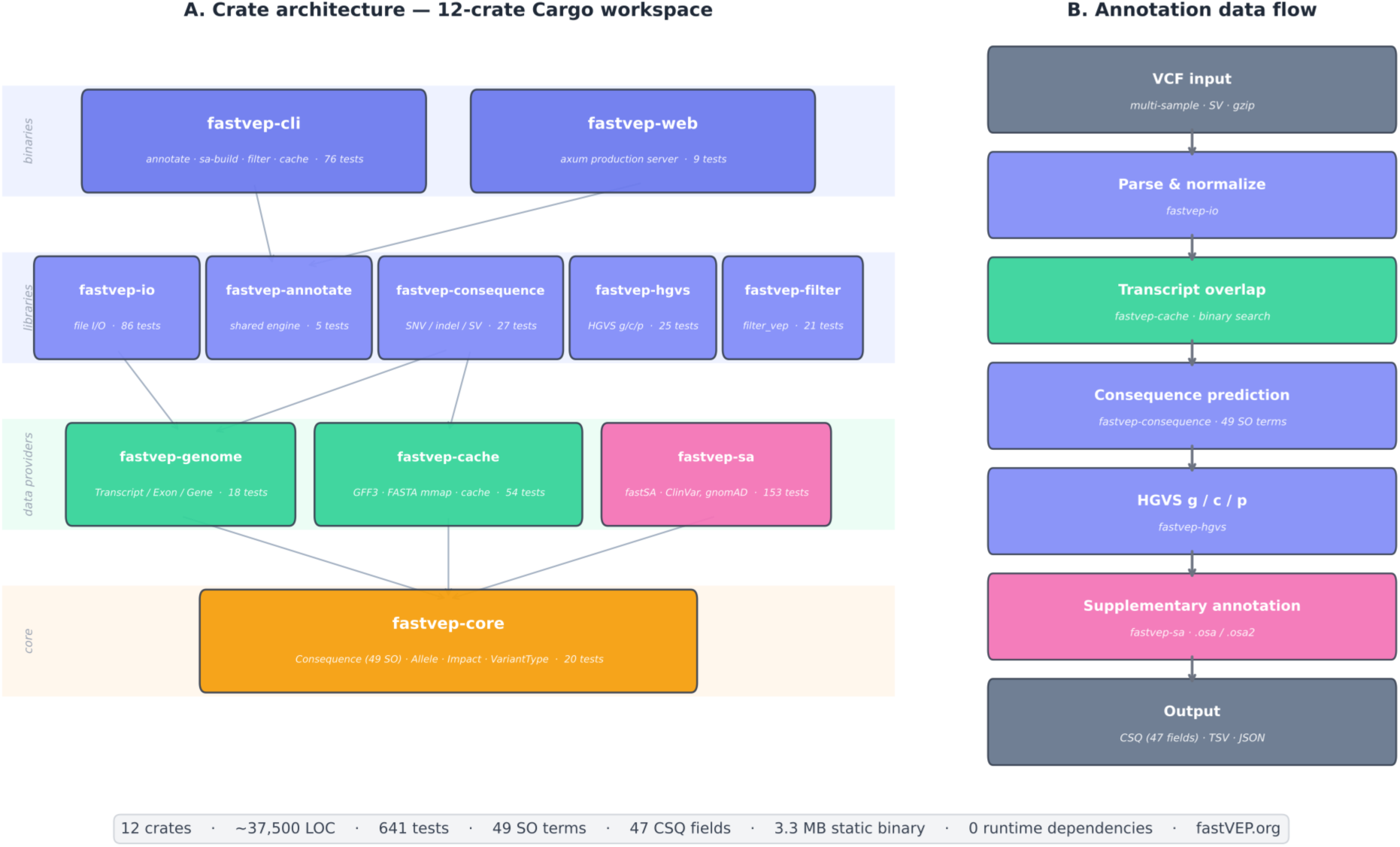
fastVEP architecture. The twelve-crate workspace, showing data flow from VCF input through consequence prediction to output formatting. Arrows indicate crate dependencies. The fastvep-consequence crate is the computational core, consuming transcript models from fastvep-cache and variant representations from fastvep-io. The fastvep-sa crate provides supplementary annotation with direct database integration (ClinVar, gnomAD, dbSNP, COSMIC, prediction scores, gene-level annotations).

**Table 1.**
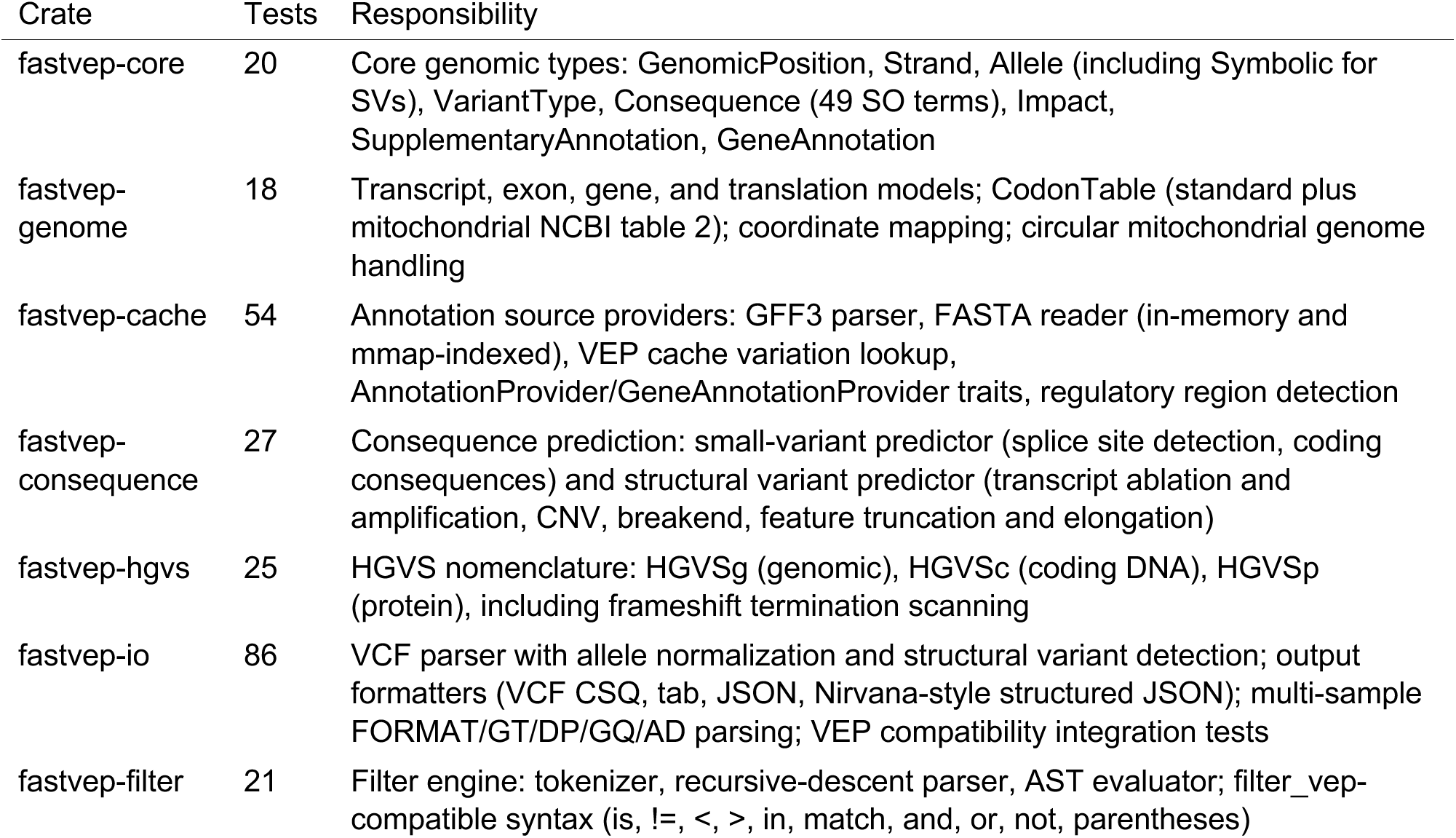
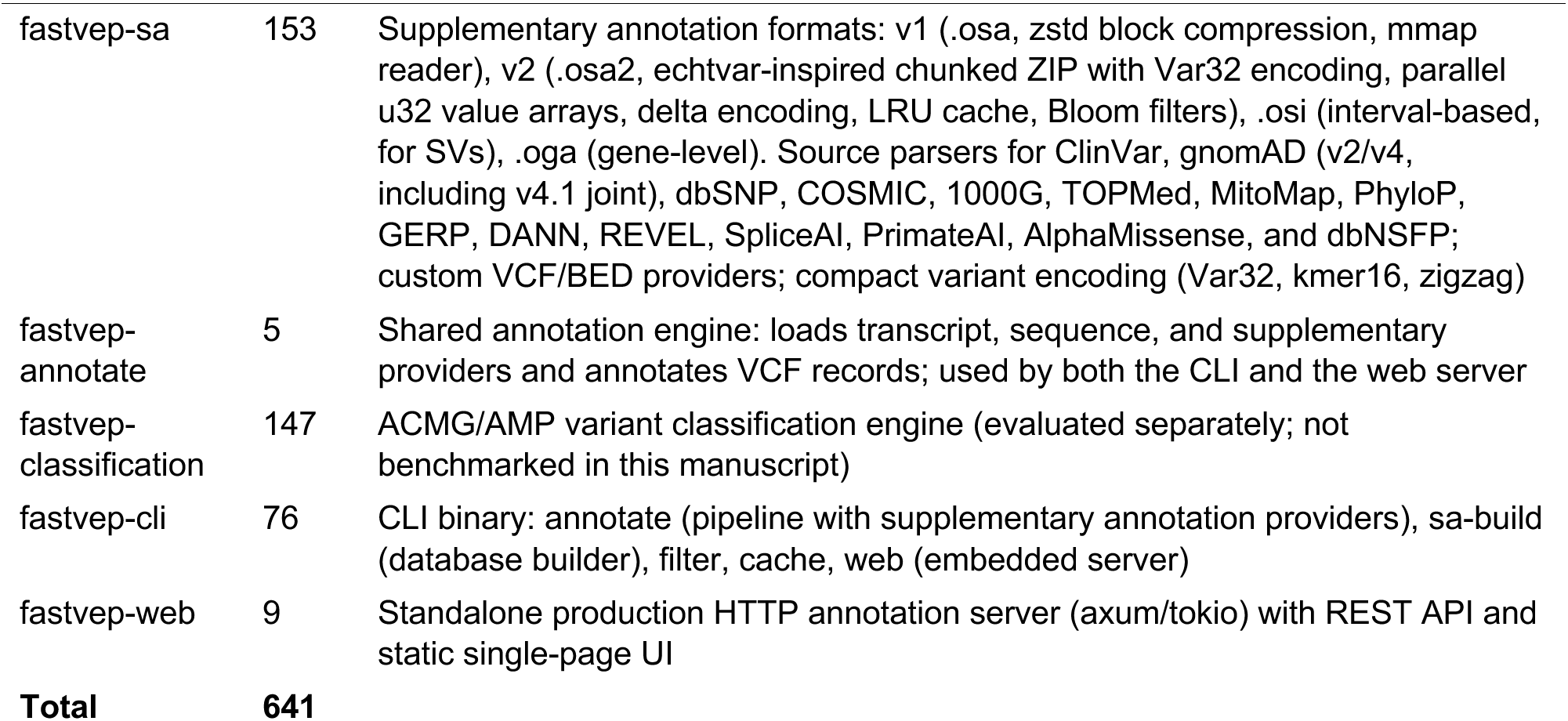
fastVEP crate architecture and responsibilities.

| Crate | Tests | Responsibility |
| --- | --- | --- |
| fastvep-core | 20 | Core genomic types: GenomicPosition, Strand, Allele (including Symbolic for SVs), VariantType, Consequence (49 SO terms), Impact, SupplementaryAnnotation, GeneAnnotation |
| fastvep-genome | 18 | Transcript, exon, gene, and translation models; CodonTable (standard plus mitochondrial NCBI table 2); coordinate mapping; circular mitochondrial genome handling |
| fastvep-cache | 54 | Annotation source providers: GFF3 parser, FASTA reader (in-memory and mmap-indexed), VEP cache variation lookup, AnnotationProvider/GeneAnnotationProvider traits, regulatory region detection |
| fastvep-consequence | 27 | Consequence prediction: small-variant predictor (splice site detection, coding consequences) and structural variant predictor (transcript ablation and amplification, CNV, breakend, feature truncation and elongation) |
| fastvep-hgvs | 25 | HGVS nomenclature: HGVSg (genomic), HGVSc (coding DNA), HGVSp (protein), including frameshift termination scanning |
| fastvep-io | 86 | VCF parser with allele normalization and structural variant detection; output formatters (VCF CSQ, tab, JSON, Nirvana-style structured JSON); multi-sample FORMAT/GT/DP/GQ/AD parsing; VEP compatibility integration tests |
| fastvep-filter | 21 | Filter engine: tokenizer, recursive-descent parser, AST evaluator; filter_vep-compatible syntax (is, !=, <, >, in, match, and, or, not, parentheses) |
| fastvep-sa | 153 | Supplementary annotation formats: v1 (.osa, zstd block compression, mmap reader), v2 (.osa2, echtvar-inspired chunked ZIP with Var32 encoding, parallel u32 value arrays, delta encoding, LRU cache, Bloom filters), .osi (interval-based, for SVs), .oga (gene-level). Source parsers for ClinVar, gnomAD (v2/v4, including v4.1 joint), dbSNP, COSMIC, 1000G, TOPMed, MitoMap, PhyloP, GERP, DANN, REVEL, SpliceAI, PrimateAI, AlphaMissense, and dbNSFP; custom VCF/BED providers; compact variant encoding (Var32, kmer16, zigzag) |
| fastvep-annotate | 5 | Shared annotation engine: loads transcript, sequence, and supplementary providers and annotates VCF records; used by both the CLI and the web server |
| fastvep-classification | 147 | ACMG/AMP variant classification engine (evaluated separately; not benchmarked in this manuscript) |
| fastvep-cli | 76 | CLI binary: annotate (pipeline with supplementary annotation providers), sa-build (database builder), filter, cache, web (embedded server) |
| fastvep-web | 9 | Standalone production HTTP annotation server (axum/tokio) with REST API and static single-page UI |
| **Total** | **641** |  |

### 2.2 Data Model

The core data model lives in fastvep-core and fastvep-genome. Genomic positions use 1-based, inclusive coordinates, matching the Ensembl convention. Alleles are represented as an enum that distinguishes nucleotide sequences, deletions, missing alleles (*), and symbolic alleles for structural variants (<DEL>, <DUP>, <INV>, and others).

The VariantType enum classifies variants as SNV, insertion, deletion, indel, MNV, or one of seven structural variant types: CopyNumberVariation, CopyNumberLoss, CopyNumberGain, TandemDuplication, Inversion, TranslocationBreakend, and ShortTandemRepeatVariation. The Consequence enum encodes 49 Sequence Ontology terms: 41 from Ensembl VEP plus 8 SV-specific terms (copy_number_change, copy_number_increase, copy_number_decrease, short_tandem_repeat_change/expansion/contraction, unidirectional_gene_fusion, and transcript_variant). Each term carries a severity rank and an impact class (HIGH, MODERATE, LOW, MODIFIER).

Transcript models capture full gene structure, including exons, coding regions, UTRs, and translation metadata. The coordinate mapping system converts in both directions between genomic, cDNA, CDS, and protein positions, and handles forward- and reverse-strand genes correctly.

### 2.3 VCF Parsing and Allele Normalization

The VCF parser handles VCF 4.1 and later, with full support for multi-allelic records, structural variant notation, star alleles, and chromosome prefix variants (chr1 versus 1). Allele normalization follows Ensembl VEP’s behavior precisely:

1. **Single-allele indels.** When the reference and alternative allele share a common first base, as VCF convention requires for indels, that base is stripped and the start position is incremented. VCF POS=100 REF=ATCG ALT=A becomes Ensembl-format start=101 ref=TCG alt=-.
2. **Multi-allelic indels.** The shared first base is stripped only when *all* non-star alleles share the same first base with the reference.
3. **Star alleles.** Preserved unchanged.

Normalization was validated with 30 integration tests derived from Ensembl VEP test suite patterns, including the test VCF files distributed with VEP release 115.

### 2.4 Consequence Prediction Engine

The consequence prediction engine is the computational core of fastVEP, implemented in fastvep-consequence (1,633 lines). For each variant, the predictor:

1. **Identifies overlapping transcripts** within a configurable upstream and downstream distance (default 5,000 bp).
2. **Maps the variant to transcript coordinates** through the genomic → cDNA → CDS → protein coordinate mapping system.
3. **Evaluates splice site proximity** across six categories:

– *Splice donor:* first 2 intronic bases at the 5’ intron boundary
– *Splice acceptor:* last 2 intronic bases at the 3’ intron boundary
– *Splice donor 5th base:* position 5 in the intron from the donor
– *Splice donor region:* positions 3-6 in the intron from the donor
– *Splice polypyrimidine tract:* 3-15 bases from the acceptor
– *Splice region:* 3-8 intronic bases or 1-3 exonic bases from any junction
4. **Determines coding consequences** by:

– Mapping the variant to the affected codons using the translateable sequence
– Translating reference and variant codons with the NCBI standard genetic code
– Classifying the amino acid change as synonymous, missense, nonsense (stop gained), stop lost, start lost, or start retained
– Separating frameshift from in-frame indels by whether the length change is divisible by three
– Scanning the altered reading frame, for frameshifts, to locate the new termination codon
5. **Handles non-coding transcripts** separately, bypassing the coding pathway entirely. Exonic positions receive non_coding_transcript_exon_variant; intronic positions receive non_coding_transcript_variant, combined with intron_variant where applicable.
6. **Appends the NMD_transcript_variant modifier** to any transcript with the nonsense_mediated_decay biotype, whatever the primary consequence. The modifier is added after all other terms and before deduplication, matching how Ensembl VEP flags variants in transcripts predicted to undergo nonsense-mediated mRNA decay.
7. **Assigns the most severe consequence** across all transcripts for summary reporting.

### 2.5 HGVS Nomenclature

fastVEP generates HGVS nomenclature following Human Genome Variation Society guidelines (den Dunnen et al., 2016):

- **HGVSg** (genomic): chr1:g.12345A>G, chr1:g.100_102del, chr1:g.100_101insTCG
- **HGVSc** (coding DNA): ENST00000001:c.51A>G, ENST00000001:c.4_6del
- **HGVSp** (protein): ENSP00000001:p.Arg41Lys, ENSP00000001:p.Arg41Ter, ENSP00000001:p.Ala527ProfsTer48

Supported notation covers substitutions, deletions, insertions, deletion-insertions (delins), duplications, frameshifts with termination position, stop-lost extensions (p.Ter100ext*?), and synonymous variants (p.Arg41=).

Frameshifts need extra work. The hgvsp_frameshift function builds both the reference and alternative translateable sequences from the CDS start through the 3’ UTR, applies the indel, and translates both in parallel. It then finds the first amino acid position where the two peptides diverge and scans downstream in the alternative peptide for the new termination codon. The resulting notation, for example p.Ala498ProfsTer28, gives the first changed residue (Ala498), the new amino acid at that position (Pro), and the distance to the new stop codon (Ter28). When the frameshifted sequence contains unresolved regions, such as incomplete 3’ UTR coverage, the function reports fsTer? for an indeterminate termination position.

### 2.6 Output Formats

fastVEP writes three output formats:

1. **VCF.** Adds a CSQ INFO field with 47 pipe-delimited annotation fields matching Ensembl VEP’s extended format: Allele, Consequence, IMPACT, SYMBOL, Gene, Feature_type, Feature, BIOTYPE, EXON, INTRON, HGVSc, HGVSp, cDNA_position, CDS_position, Protein_position, Amino_acids, Codons, Existing_variation, REF_ALLELE, UPLOADED_ALLELE, DISTANCE, STRAND, FLAGS, CANONICAL, SYMBOL_SOURCE, HGNC_ID, MANE, MANE_SELECT, MANE_PLUS_CLINICAL, TSL, APPRIS, CCDS, ENSP, SOURCE, HGVS_OFFSET, SIFT, PolyPhen, AF, CLIN_SIG, SOMATIC, PHENO, PUBMED, MOTIF_NAME, MOTIF_POS, HIGH_INF_POS, MOTIF_SCORE_CHANGE, and TRANSCRIPTION_FACTORS. Special characters are escaped as VEP does: commas → &, semicolons → %3B, pipes → &.
2. **Tab-delimited.** One line per variant-allele-transcript combination, with the 17 default columns of VEP’s default output.
3. **JSON.** A structured representation with nested transcript consequence arrays, suited to programmatic consumption.

### 2.7 Annotation Source Architecture

Annotation sources are accessed through a trait-based provider architecture:

- **TranscriptProvider** returns transcripts overlapping a genomic region. The IndexedTranscriptProvider implementation groups transcripts by chromosome into sorted arrays and uses binary search (partition_point) for O(log n + k) overlap queries, where n is the number of transcripts on the chromosome and k is the number of overlapping results. This approach, inspired by echtvar’s chunked binary search strategy (Pedersen, 2023), replaces naive linear scans and is what makes large gene models tractable (for example, 11,605 transcripts on chromosome 22).
- **SequenceProvider** supplies reference sequences for regions. FastaSequenceProvider uses memory-mapped FASTA with .fai index files for zero-copy reference access through the memmap2 crate.
- **VariationProvider** looks up co-located known variants in tabix-indexed VEP cache files (all_vars.gz per chromosome). The TabixVariationProvider implementation matches alleles rather than positions alone, accounting for multi-allelic sites, strand orientation (with automatic reverse-complement matching for minus-strand records), and VCF-to-Ensembl deletion marker normalization. For each match, it extracts the dbSNP identifier, allele-specific population frequencies (gnomAD and 1000 Genomes continental populations), minor allele frequency, clinical significance, somatic status, phenotype associations, and PubMed references. These populate the Existing_variation, AF, CLIN_SIG, SOMATIC, PHENO, and PUBMED CSQ fields.

Four optimizations cut startup and annotation latency:

1. **O(T+E) indexed GFF3 assembly.** Transcript assembly from GFF3 features runs in O(T+E) time, where T is transcripts and E is exons, replacing an earlier O(T*E) nested iteration. GFF3 loading is 12-80x faster as a result.
2. **Binary transcript cache.** Transcripts parsed from GFF3 are serialized with bincode and gzip compression, so later runs start from cache: approximately 1.4 seconds for 7K yeast transcripts and 2.6 seconds for 509K human transcripts.
3. **Tabix-indexed GFF3 loading.** When a tabix index (.tbi) accompanies the GFF3 file, fastVEP loads only transcripts overlapping the variant regions, reducing both startup time and memory use.
4. **Memory-mapped FASTA.** Reference sequences are read through memory-mapped I/O with .fai index files, giving zero-copy access without loading the genome into memory.

Because the design is trait-based, annotation sources can be swapped transparently. The GFF3 provider can be replaced with a binary cache reader, or the tabix variation provider with a database-backed implementation, without touching the prediction engine.

### 2.8 Web Interface

fastVEP includes a built-in web interface, started with fastvep web --gff3 <FILE> --fasta <FILE> --port 8080. The interface is a single-page application of roughly 1,750 lines of HTML, CSS, and JavaScript, embedded directly in the binary through Rust’s include_str! macro, so it needs no external files or dependencies.

It runs in two modes. Served by the fastVEP binary (fastvep web), annotation requests go to the Rust backend through a REST API (POST /api/annotate), with the same performance and accuracy as the CLI. Users can upload custom GFF3 files from the browser, which the Rust GFF3 parser handles server-side (POST /api/upload-gff3), avoiding slow client-side parsing of large annotation files. Deployed as a static page, for instance on GitHub Pages, the interface falls back to a client-side JavaScript annotation engine that reimplements core consequence prediction for interactive use without a server.

Interface features include:

- A VCF input area with example datasets (coding, splicing, and non-coding variants)
- Custom GFF3 upload with server-side or client-side parsing
- Configurable annotation options (upstream and downstream distance, pick mode for canonical transcripts)
- Light and dark mode with system preference detection
- An interactive results table with all annotation fields (HGVSg/c/p, canonical, strand, distance), column sorting, and search
- A summary view with bar charts of impact and consequence distributions
- Raw output in VCF (47-field CSQ), tab-delimited, or JSON format
- Automatic backend detection with transparent fallback to client-side annotation

### 2.9 Supplementary Annotation Framework (fastSA)

fastVEP includes a native supplementary annotation framework, fastSA, for direct integration with clinical and population databases. Its design is inspired by Illumina Nirvana’s NSA architecture (Stromberg et al., 2024). The framework defines three binary file formats.

#### 1. Position and allele-level annotations (.osa / .osa2)

Two format generations are supported. The v1 format (.osa) uses zstd block-compressed data files with per-chromosome indexing and memory-mapped I/O. The v2 format (.osa2), inspired by echtvar (Pedersen, 2023), uses ZIP-based archives built around:

- Genomic chunks of about 1 MB (position >> 20)
- Compact 32-bit variant encoding (Var32: 20-bit position, 2-bit ref_len, 2-bit alt_len, 8-bit encoded bases)
- Parallel u32 value arrays in place of JSON strings
- Delta encoding of variant keys
- Float quantization (value × multiplier → u32) and categorical string-to-index encoding
- Zigzag encoding for signed values
- LRU chunk caching

Together these remove JSON parsing from the hot path and allow O(log n) binary search on 4-byte keys rather than variable-length string comparisons. Bloom filters answer negative lookups quickly, skipping chunk decompression when a position is definitely not annotated.

#### 2. Interval-level annotations (.osi)

For structural variant databases such as gnomAD SV, ClinGen dosage, and DGV, where annotations cover genomic regions. Supports overlap queries with reciprocal overlap calculation.

#### 3. Gene-level annotations (.oga)

For gene-keyed databases such as OMIM and gnomAD gene constraint metrics, looked up by gene symbol.

The AnnotationProvider trait defines the interface for all supplementary sources. It supports batch preloading, which decompresses the relevant blocks before parallel annotation begins, and thread-safe parallel querying through Rust’s Send + Sync constraints. The sa-build CLI subcommand builds fastSA files from standard source formats:

- **Clinical databases:** ClinVar (VCF: CLNSIG, CLNREVSTAT, CLNDN), dbSNP (VCF: RS IDs, global MAF), COSMIC (VCF: COSV IDs, gene, sample counts)
- **Population frequencies:** gnomAD (VCF: AF per population; v2 exomes and genomes through v4.0 and v4.1, including v4.1 joint frequencies), 1000 Genomes (VCF: 5 continental populations), TOPMed (VCF: AF, AC, AN), MitoMap (TSV: disease, status)
- **Prediction scores:** PhyloP (wigFix), GERP (TSV), DANN (TSV), REVEL (CSV), SpliceAI (VCF: delta scores), PrimateAI (TSV), AlphaMissense (TSV: pathogenicity score and class), dbNSFP (TSV: SIFT/PolyPhen)
- **Gene-level:** OMIM genemap2 (TSV: MIM numbers, phenotypes), gnomAD gene constraint (TSV: pLI, LOEUF, mis_z, syn_z)

Under the default --format auto, sa-build writes v2 .osa2 for the allele- and position-level sources that support it and v1 .osa otherwise. Both formats are queried transparently at annotation time.

### 2.10 Structural Variant Support

fastVEP annotates structural variants including deletions (<DEL>), duplications (<DUP>, <DUP:TANDEM>), inversions (<INV>), copy number variants (<CNV>, <CN0>-<CN4>), breakends (<BND>), insertions (<ins>), and short tandem repeats (<STR>). The VCF parser detects symbolic alleles, reads SV-specific INFO fields (SVTYPE, END, SVLEN, CIPOS, CIEND, MATEID), and classifies each variant into a VariantType category.

The SvConsequencePredictor assigns consequences from the overlap between the SV interval and transcript models:

- **Complete containment:** transcript_ablation (deletions) or transcript_amplification (duplications)
- **Partial overlap:** feature_truncation, feature_elongation, coding_sequence_variant (if the CDS is affected), splice_acceptor_variant (if splice sites are affected)
- **Copy number:** copy_number_change, copy_number_increase, copy_number_decrease
- **Breakends:** transcript_variant, coding_sequence_variant (if landing in the CDS)
- **Repeat expansions:** short_tandem_repeat_change

### 2.11 Filter Engine

fastVEP implements a filter engine compatible with Ensembl VEP’s filter_vep tool. It has three parts: a tokenizer that handles field names, comparison operators (is, !=, <, >, <=, >=, in, match), logical operators (and, or, not), and parenthesized grouping; a recursive-descent parser that produces a typed AST; and an evaluator that resolves field values from annotation output and applies the filter logic. String comparisons are case-insensitive, numeric comparisons parse field values to f64, and the in operator understands ampersand-separated VEP consequence fields.

### 2.12 Multi-Sample and Regulatory Support

fastVEP parses VCF FORMAT and sample columns to extract per-sample genotype (GT), depth (DP), genotype quality (GQ), and allele depth (AD). Genotypes are classified as het, hom_ref, hom_alt, or missing, which supports trio-based filtering such as de novo variant detection.

Regulatory annotations come from Ensembl regulatory build GFF3 files, identifying promoters, enhancers, CTCF binding sites, TF binding sites, and open chromatin regions. Mitochondrial variants are handled with circular coordinate wrapping and the vertebrate mitochondrial codon table (NCBI translation table 2), which differs from the standard table at AGA/AGG (Stop instead of Arg), ATA (Met instead of Ile), and TGA (Trp instead of Stop).

**Table 2.** Single-threaded scaling on GIAB HG002 chromosome 22. Apple M4, median of 3 runs, full Ensembl GRCh38 release 115 annotations with FASTA reference and HGVS generation. fastVEP used its binary transcript cache; VEP ran in indexed --gff mode.

| Variants | fastVEP | Ensembl VEP | Speedup |
| --- | --- | --- | --- |
| 1,000 | 2.68s | 0.45s | 0.17x (VEP faster) |
| 5,000 | 2.86s | 6.29s | 2.2x |
| 10,000 | 3.25s | 13.41s | 4.1x |
| 50,284 | 5.09s | 93.95s | 18.5x |

### 2.13 Why Rust

Rust was chosen for four properties that matter in bioinformatics tool development:

1. **Zero-cost abstractions.** Trait-based polymorphism compiles to direct function calls, with no runtime dispatch overhead.
2. **Memory safety without garbage collection.** The ownership system rules out whole classes of bugs, including use-after-free, data races, and null pointer dereferences, without the unpredictable pauses of garbage-collected languages.
3. **Static binary compilation.** fastVEP compiles to a single 3.3 MB binary with no runtime dependencies, which simplifies deployment on HPC systems, in Docker containers, and in clinical pipelines.
4. **Ecosystem maturity.** The Rust bioinformatics ecosystem offers production-quality crates for genomic file formats (noodles, rust-htslib), interval trees (coitrees), and parallelism (rayon).

## 3. Results

### 3.1 Single-Threaded Scaling on Chromosome 22

We benchmarked fastVEP against Ensembl VEP v115.1 on the GIAB HG002 v4.2.1 benchmark VCF (4,048,342 high-confidence variants). Both tools received identical inputs: the full Ensembl GRCh38 release 115 GFF3 gene model (508,530 transcripts), the GRCh38 primary-assembly FASTA reference, and equivalent options (--hgvs --symbol --canonical). To isolate raw engine efficiency, both were run single-threaded (Tables 2-3, **Figure 2**). fastVEP used its binary transcript cache; Ensembl VEP ran in Docker (ensemblorg/ensembl-vep:release_115.1) in indexed --gff mode against a bgzip- and tabix-indexed GFF3.

**Figure 2.**
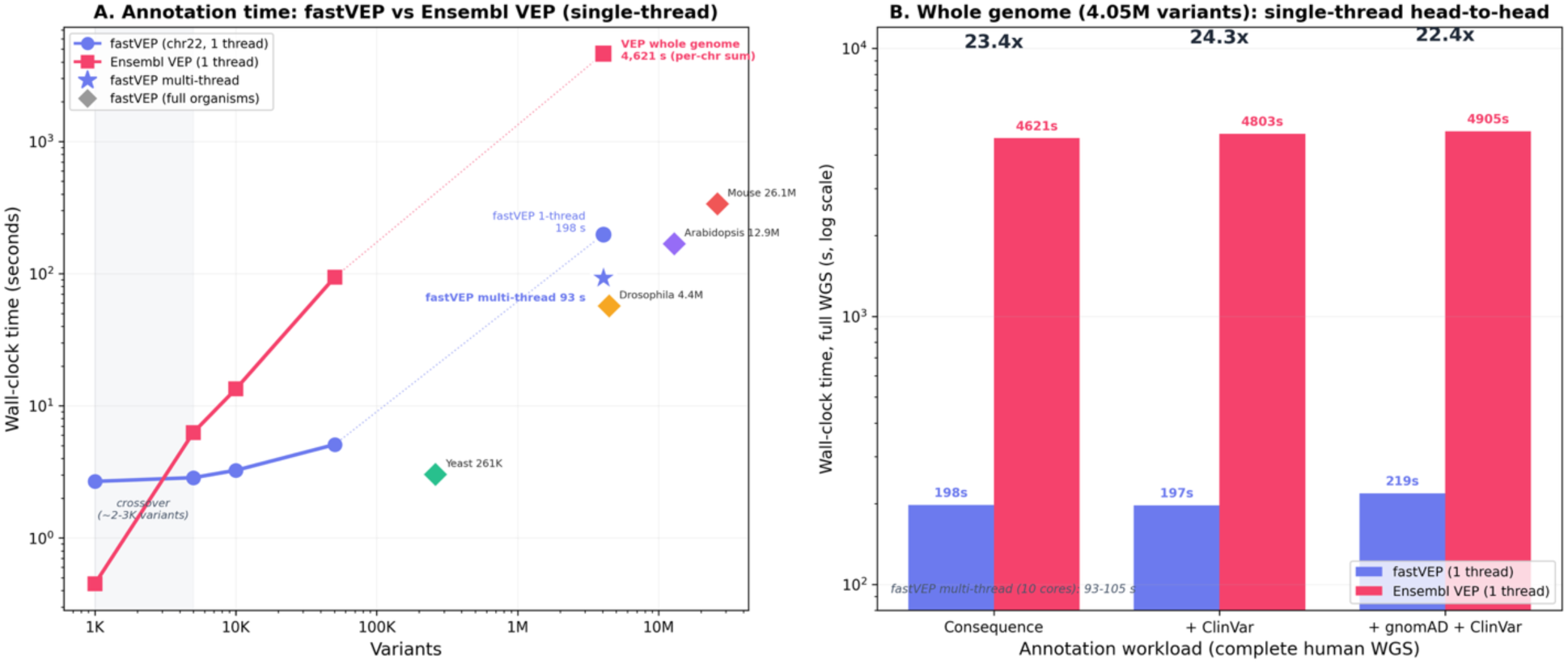
fastVEP versus Ensembl VEP: single-threaded performance. (A) Wall-clock annotation time (log-log) for both tools on the chromosome 22 scaling series (1K-50K GIAB HG002 variants, connected markers), together with fastVEP multi-organism results (diamonds: yeast 260K, Drosophila 4.4M, Arabidopsis 12.9M, mouse 26M) and the three directly measured whole-genome points at 4.05M variants (Ensembl VEP per-chromosome sum, 4,621 s; fastVEP single-thread, 198 s; fastVEP multi-thread, 93 s, star). fastVEP carries a fixed 2.6 s binary-cache load, so the two tools cross over between 1,000 and 5,000 variants (shaded), after which fastVEP dominates. (B) Whole-genome single-threaded head-to-head across the three annotation workloads (consequence-only, +ClinVar, +gnomAD+ClinVar) on a log time axis, with per-workload speedups (22.4-24.3x); fastVEP’s default multi-threaded time (86 s) applies to all three workloads.

**Table 3.**
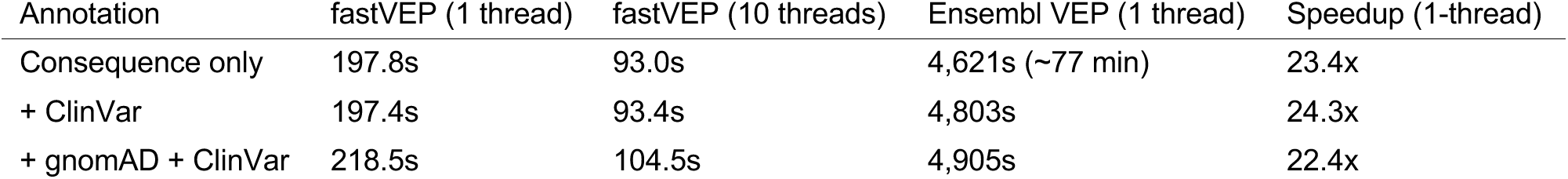
Head-to-head on the complete GIAB HG002 WGS (4,048,342 variants, GRCh38, 508,530 transcripts). Both tools measured on the same Apple M4 system. Supplementary annotations were supplied to fastVEP as native fastSA databases and to Ensembl VEP as --custom tabix-indexed VCFs built from identical source data. Ensembl VEP full-genome times are per-chromosome sums (minimum of two runs). fastVEP times are shown single-threaded (RAYON_NUM_THREADS=1) and multi-threaded (default, 10 cores; best of three runs); the speedup column is the like-for-like single-threaded comparison.

| Annotation | fastVEP (1 thread) | fastVEP (10 threads) | Ensembl VEP (1 thread) | Speedup (1-thread) |
| --- | --- | --- | --- | --- |
| Consequence only | 197.8s | 93.0s | 4,621s (~77 min) | 23.4x |
| + ClinVar | 197.4s | 93.4s | 4,803s | 24.3x |
| + gnomAD + ClinVar | 218.5s | 104.5s | 4,905s | 22.4x |

We first measured how the comparison scales with input size on chromosome 22 (Table 2). fastVEP loads its whole-genome binary transcript cache up front, which costs about 2.6 seconds, whereas VEP lazily fetches only the tabix-indexed GFF regions overlapping the input. VEP is therefore faster on very small inputs. The two tools cross over between 1,000 and 5,000 variants, after which fastVEP’s per-variant throughput dominates, reaching 18.5x at roughly 50,000 variants.

### 3.2 Clinical Whole-Genome Head-to-Head

At genome scale the fixed startup cost is fully amortized, and fastVEP’s advantage is largest. On the complete GIAB HG002 WGS, fastVEP is 22-24x faster than single-threaded Ensembl VEP across consequence-only and annotation-augmented workloads (Table 3).

A single-process whole-genome VEP run exhausts the default container memory above roughly 2.7 million variants, because transcript structures accumulate across the genome as annotation proceeds. VEP’s genome-wide time was therefore measured per chromosome and summed. This also caps its resident memory at a single chromosome and represents its best case. Each per-chromosome time is the minimum of two runs, which removes transient system contention (Methods).

With its default multi-threaded pipeline on 10 cores, fastVEP annotates the same complete WGS in 93 seconds, or 43,500 variants per second. That is about 2.1x its single-threaded speed and consistent with the 86 seconds recorded for this genome in the multi-organism benchmark (Table 7). A complete clinical WGS therefore takes under two minutes, fast enough for routine use in clinical sequencing pipelines, where annotation has traditionally been a multi-hour step. The practical gap is widest exactly where population- and clinical-scale work needs it most: the complete human genome.

These whole-genome VEP times are **directly measured**, with each chromosome run separately and the wall-clock times summed, rather than extrapolated. They supersede an earlier version of this benchmark that projected VEP’s full-genome runtime from chromosome 22 throughput and reported a substantially larger nominal speedup. That projection assumed a slower per-variant rate and did not correspond to a like-for-like measured run on an identically indexed gene model. In the present comparison, both tools read the **same bgzip- and tabix-indexed** GRCh38 release 115 GFF3 at every scale. VEP’s full-genome result was obtained the only way it completes within the available memory, per chromosome and then summed. The resulting 22-24x single-threaded advantage for fastVEP is therefore conservative and reproducible.

### 3.3 Annotation Accuracy Against Ensembl VEP

Accuracy was validated by field-level comparison against Ensembl VEP release 115.1 on 173 real human chromosome 22 variants from the VEP example dataset (**Figure 3**). Both tools ran with the same GFF3 annotations and GRCh38 FASTA reference and equivalent flags (--hgvs --symbol --canonical).

**Figure 3.**
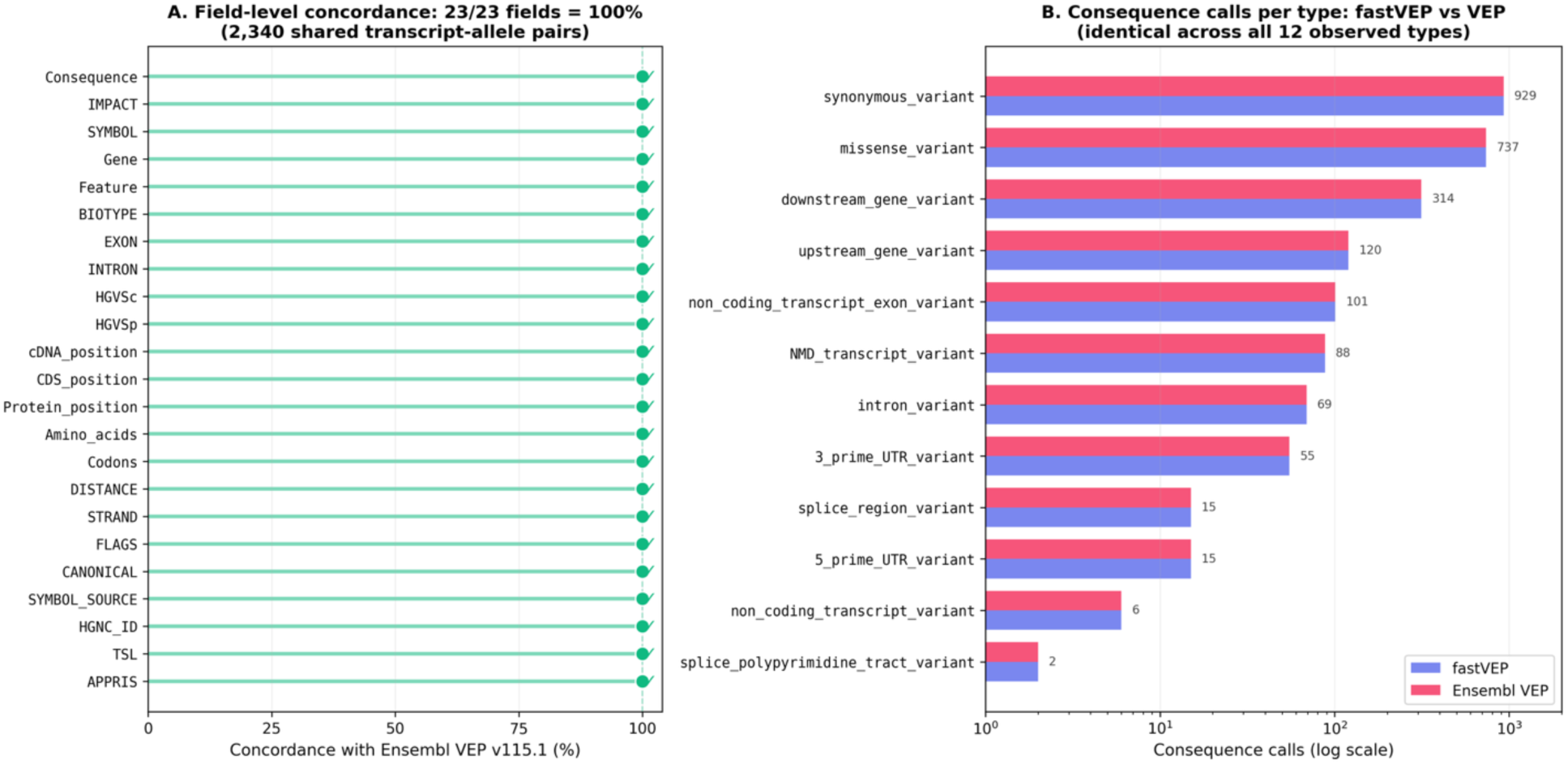
VEP concordance. (A) Field-level accuracy of fastVEP against Ensembl VEP v115.1 on 2,340 shared transcript-allele pairs from 173 chromosome 22 variants: all 23 compared annotation fields match. (B) Consequence calls per Sequence Ontology type (log scale) for fastVEP and Ensembl VEP on the same set; the two tools produce identical call counts across all 12 observed consequence types.

All 23 compared annotation fields matched on all 2,340 shared (allele, transcript) pairs (Figure 3A). fastVEP annotated 35 additional lncRNA transcripts absent from the VEP output, a consequence of different default transcript filtering, and missed none. The consequence type distribution was identical across all 12 consequence types observed in the dataset: missense_variant (737), synonymous_variant (929), intron_variant (69), upstream_gene_variant (120), downstream_gene_variant (314), 3_prime_UTR_variant (55), 5_prime_UTR_variant (15), NMD_transcript_variant (88), non_coding_transcript_exon_variant (101), non_coding_transcript_variant (6), splice_region_variant (15), and splice_polypyrimidine_tract_variant (2).

### 3.4 Resource Efficiency

**Table 4.** Resource usage comparison between fastVEP and Ensembl VEP.

| Metric | Ensembl VEP (Perl) | fastVEP (Rust) | Improvement |
| --- | --- | --- | --- |
| Binary/install size | ~200 MB (with dependencies) | 3.3 MB | 63x smaller |
| Startup (7K transcripts, yeast) | 5-15 sec | 1.4 sec | 4-11x faster |
| Startup (509K transcripts, human) | N/A (no binary cache) | 2.6 sec | — |
| External dependencies | Perl 5.22+, DBI, 10+ CPAN modules | None | — |
| Installation | INSTALL.pl + compile C extensions | cargo build --release | — |

The binary transcript cache (bincode with gzip compression) keeps startup fast even for the largest genomes: about 1.4 seconds for yeast (7K transcripts) and 2.6 seconds for the full human genome (508K transcripts), loaded straight from the serialized cache without reparsing GFF3. The release binary is compiled with link-time optimization and a single codegen unit, yielding a 3.3 MB statically linked executable. Variants are processed as a stream rather than buffered in memory, and memory-mapped FASTA access avoids loading the reference genome.

### 3.5 Output Format Performance

**Table 5.** Output format comparison for 10,000 GIAB HG002 chromosome 22 variants against full GRCh38 (508,530 transcripts), single-threaded (median of 3 runs).

| Format | Time (sec) | Output size |
| --- | --- | --- |
| VCF (CSQ, 47 fields) | 3.12 | 46,511 KB |
| Tab-delimited | 2.97 | 25,312 KB |
| JSON | 3.12 | 83,507 KB |

All three formats perform much the same: the roughly 2.6-second binary-cache startup dominates, and per-variant formatting overhead is negligible. JSON files are the largest, about 3.3x the size of tab-delimited output, because the structured representation is verbose, but they give the richest programmatic access to nested consequence data.

### 3.6 Supplementary Annotation Format (fastSA v2)

The v2 .osa2 format is the default output of sa-build for supported sources. To evaluate it on real databases, we built v1 .osa and v2 .osa2 representations of ClinVar and REVEL from their standard source files and compared on-disk size, build time, and annotation output (Table 6).

**Table 6.** fastSA v1 versus v2 on real annotation databases (GRCh38).

| Source | Records | v1 (.osa) | v2 (.osa2) | Ratio | Build v1 | Build v2 |
| --- | --- | --- | --- | --- | --- | --- |
| ClinVar | 4.44M | 48.1 MB | 38.9 MB | 0.81x | 19.0s | 24.2s |
| REVEL (chr1) | 8.25M | 40.5 MB | 19.0 MB | 0.47x | 9.7s | 10.3s |

The v2 format stores both databases more compactly, at 0.81x (ClinVar) and 0.47x (REVEL) of the v1 size, for a modest increase in build time. The annotation output is byte-identical between formats: annotating a 673,000-variant real call set against ClinVar produced md5-identical VCFs under v1 (15.0 s) and v2 (14.3 s). The upgrade is therefore a pure storage and lookup optimization with no effect on annotation content, so databases can be migrated to .osa2 without revalidation.

### 3.7 Cross-Organism Annotation

To test generality, we benchmarked annotation across five model organisms using full Ensembl GFF3 gene models with FASTA reference and HGVS generation (Table 7, **Figure 4**). All runs used the binary transcript cache for startup and report the median of 3 runs.

**Figure 4.**
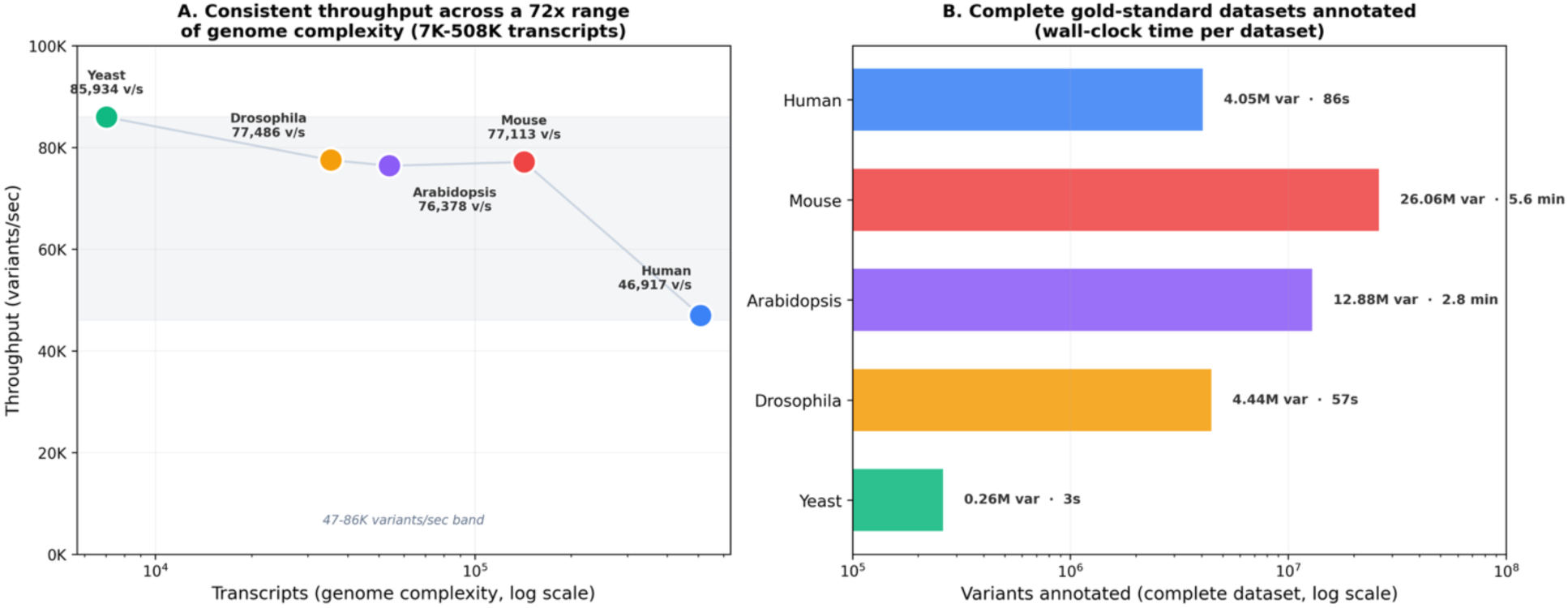
Multi-organism throughput and scale. (A) Annotation throughput (variants per second) versus genome complexity (transcripts, log scale) for five organisms, using complete gold-standard VCFs with full Ensembl genome annotations; throughput stays within a 47K-86K variants per second band across a 72x range of genome complexity (7K-508K transcripts). (B) Absolute scale of the complete datasets annotated (variants, log scale), with wall-clock time per dataset, from 0.26M yeast variants in 3 s to 26.06M mouse variants in 5.6 minutes.

**Table 7.** Cross-organism annotation performance using full Ensembl genome annotations with gold-standard variant call sets. All benchmarks use full Ensembl release 115 GFF3 gene models, FASTA reference with memory-mapped access, and HGVS generation. Variant sources: Ensembl/SGD variation (yeast), DGRP2 (Drosophila), 1001 Genomes (Arabidopsis), Ensembl/EVA variation (mouse), GIAB HG002 v4.2.1 (human). Median of 3 runs.

| Organism | Assembly | Transcripts | Variants | Source | Time (sec) | Throughput (v/s) |
| --- | --- | --- | --- | --- | --- | --- |
| Yeast (*S. cerevisiae*) | R64-1-1 | 7,036 | 260,526 | Ensembl/SGD | 3.03 | 85,934 |
| Drosophila (*D. melanogaster*) | BDGP6.54 | 35,442 | 4,438,427 | DGRP2 | 57.28 | 77,486 |
| Arabidopsis (*A. thaliana*) | TAIR10 | 54,013 | 12,883,854 | 1001 Genomes | 168.68 | 76,378 |
| Mouse (*M. musculus*) full | GRCm39 | 142,626 | 26,062,054 | MGP CAST/EiJ | 337.97 | 77,113 |
| Human (*H. sapiens*) full WGS | GRCh38 | 508,530 | 4,048,342 | GIAB HG002 | 86.29 | 46,917 |

fastVEP annotated complete gold-standard datasets for all five organisms at 47,000 to 86,000 variants per second. The mouse benchmark uses the CAST/EiJ strain from the Mouse Genomes Project, the most genetically divergent common laboratory strain, with 26 million SNPs and indels against 142,626 transcripts; it finished in 5.6 minutes at 77K v/s. The Arabidopsis 1001 Genomes dataset, 12.9M variants from 1,135 accessions, finished in 2.8 minutes at 76K v/s. The clinical benchmark, a complete human WGS of 4.05M GIAB HG002 high-confidence variants against 508,530 transcripts, finished in 86 seconds. C. elegans was excluded because the CaeNDR gold-standard VCF source is currently unavailable.

### 3.8 Consequence Prediction on a Complete Genome

We applied fastVEP to the complete GIAB HG002 WGS (4,048,342 variants) against the full Ensembl GRCh38 gene model (508,530 transcripts). The run produced 50,109,169 consequence calls across all overlapping transcripts, spanning 26 distinct SO terms (Table 8, **Figure 5**).

**Figure 5.**
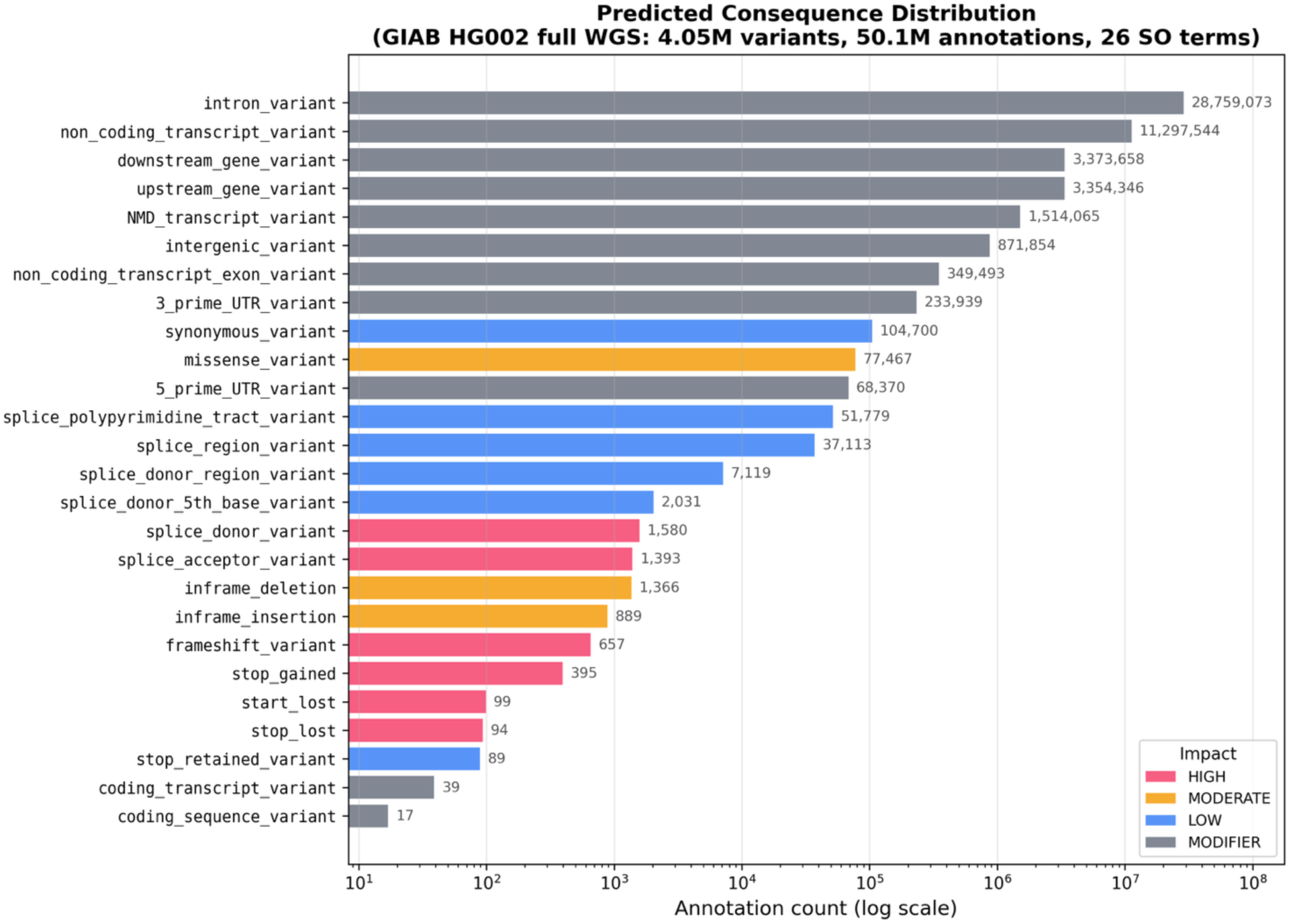
Consequence distribution. Distribution of 50.1 million predicted consequences from the complete GIAB HG002 WGS (4,048,342 variants) annotated against the full Ensembl GRCh38 gene model (508,530 transcripts). The distribution covers 26 SO consequence types, with intronic (57.4%) and non-coding transcript (22.5%) variants predominating.

**Table 8.** Genome-wide consequence distribution across the complete GIAB HG002 WGS (4,048,342 variants; 50,109,169 consequence calls). Fractions are of total consequence calls.

| Consequence | Count | Fraction |
| --- | --- | --- |
| intron_variant | 28,759,073 | 57.4% |
| non_coding_transcript_variant | 11,297,544 | 22.5% |
| downstream_gene_variant | 3,373,658 | 6.7% |
| upstream_gene_variant | 3,354,346 | 6.7% |
| NMD_transcript_variant | 1,514,065 | 3.0% |
| intergenic_variant | 871,854 | 1.7% |
| non_coding_transcript_exon_variant | 349,493 | 0.7% |
| 3_prime_UTR_variant | 233,939 | 0.5% |
| synonymous_variant | 104,700 | 0.21% |
| missense_variant | 77,467 | 0.15% |
| 5_prime_UTR_variant | 68,370 | 0.14% |
| splice_polypyrimidine_tract_variant | 51,779 | 0.10% |
| splice_region_variant | 37,113 | 0.07% |
| splice_donor_region_variant | 7,119 | 0.014% |
| splice_donor_5th_base_variant | 2,031 | 0.004% |
| splice_donor_variant | 1,580 | 0.003% |
| splice_acceptor_variant | 1,393 | 0.003% |
| inframe_deletion | 1,366 | 0.003% |
| inframe_insertion | 889 | 0.002% |
| frameshift_variant | 657 | 0.001% |
| stop_gained | 395 | <0.001% |
| start_lost | 99 | <0.001% |
| stop_lost | 94 | <0.001% |
| stop_retained_variant | 89 | <0.001% |
| coding_transcript_variant | 39 | <0.001% |
| coding_sequence_variant | 17 | <0.001% |

Intronic (57.4%) and non-coding transcript (22.5%) consequences predominate, followed by upstream and downstream gene variants at 6.7% each. High-impact coding consequences, namely missense, stop gained, frameshift, and splice donor or acceptor variants, together account for well under 0.5% of calls. This is what one expects for a single genome dominated by common non-coding variation. It also shows that fastVEP resolves the full spectrum of SO consequence classes at genome scale, including the fine-grained splice categories that many annotation tools omit: splice donor 5th base, splice donor region, and polypyrimidine tract.

### 3.9 Test Suite Coverage

Correctness was validated with a test suite of 641 tests, all passing under rustc 1.94.1. Table 9 summarizes coverage of the principal functional areas. The counts are representative rather than mutually exclusive, and further tests cover the I/O, CLI, shared annotation-engine, web-server, and classification crates.

**Table 9.**
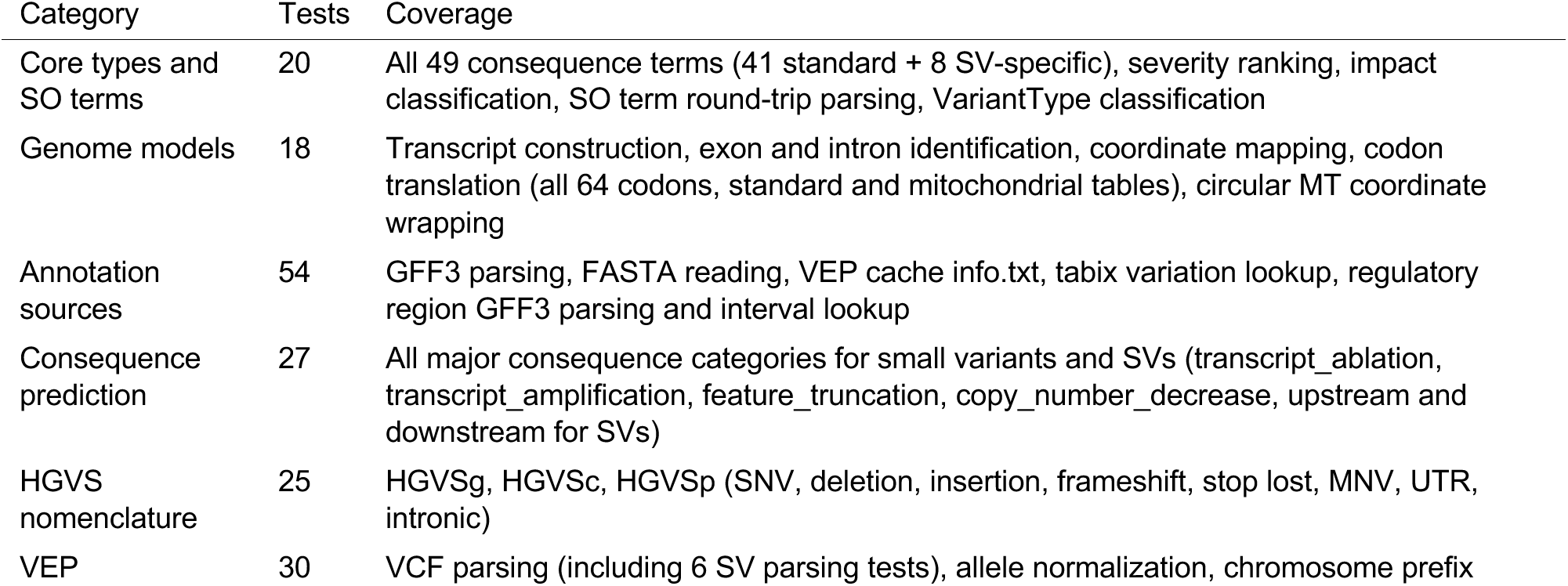
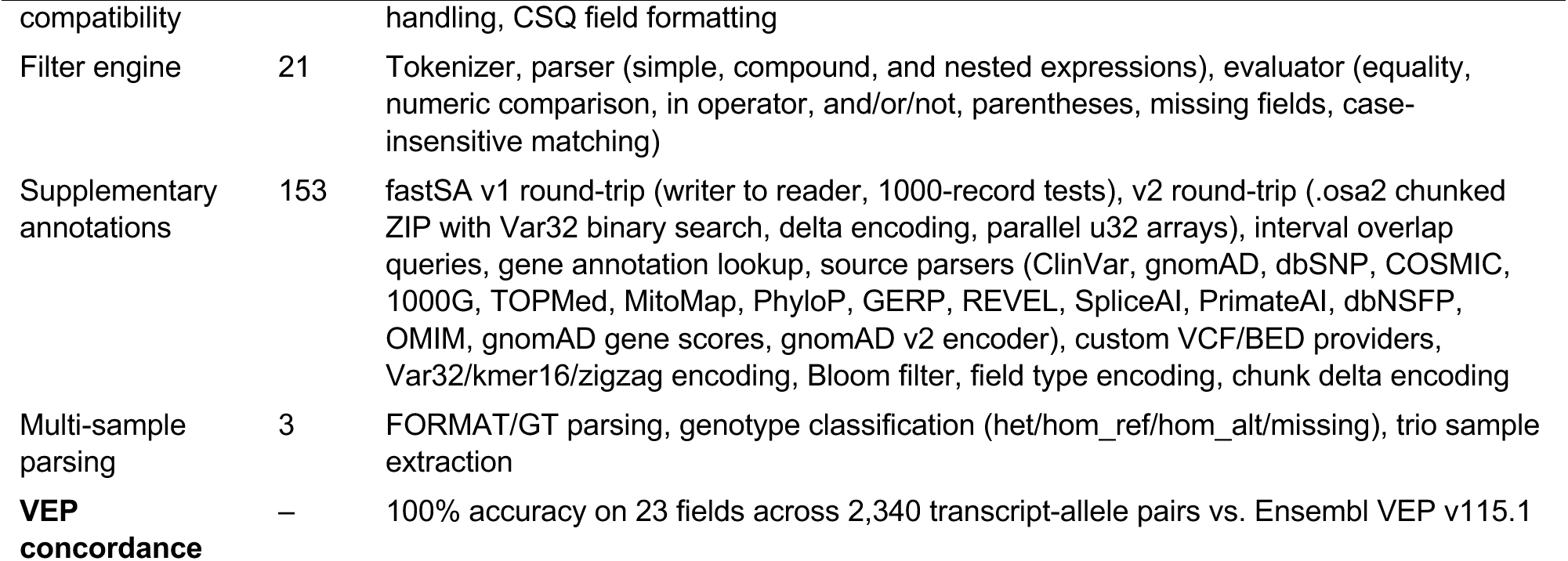
Test suite coverage.

### 3.10 Feature Comparison

**Table 10.** Feature comparison between fastVEP, Ensembl VEP, and Illumina Nirvana.

| Feature | Ensembl VEP | Nirvana | fastVEP |
| --- | --- | --- | --- |
| Language | Perl 5.22+ | C# | Rust |
| SO consequence terms | ~35 | ~40 | 49 |
| CSQ output fields | 30 (default) | N/A (JSON) | 47 (extended) |
| Splice site categories | 3 | 3 | 6 (+ 5th base, donor region, polypyrimidine) |
| VCF input | Yes | Yes | Yes |
| Structural variants | Limited | Yes | Yes (DEL, DUP, INV, CNV, BND, STR) |
| ClinVar | Via plugin | Yes | Yes (direct .osa) |
| gnomAD | Via plugin | Yes | Yes (direct .osa) |
| dbSNP | Via cache | Yes | Yes (direct .osa) |
| COSMIC | Via plugin | Yes | Yes (direct .osa) |
| 1000 Genomes | Via cache | Yes | Yes (direct .osa) |
| PhyloP/GERP | Via plugin | Yes | Yes (direct .osa) |
| REVEL/SpliceAI | Via plugin | Yes | Yes (direct .osa) |
| SIFT/PolyPhen | Yes | Yes | Yes (via dbNSFP .osa) |
| OMIM | Via plugin | Yes | Yes (gene-level .oga) |
| Gene constraint (pLI) | Via plugin | Yes | Yes (gene-level .oga) |
| Regulatory features | Yes | Yes | Yes (GFF3) |
| Filter engine | filter_vep | – | Yes (filter_vep-compatible) |
| Multi-sample genotypes | Yes | Yes | Yes (FORMAT/GT parsing) |
| Custom annotations | Yes (–custom) | Yes | Yes (VCF/BED) |
| Mitochondrial codon table | Yes | Yes | Yes (NCBI table 2) |
| JSON output (structured) | Yes | Yes (primary) | Yes (Nirvana-style + VEP-style) |
| Binary transcript cache | – | Yes | Yes (bincode+gzip) |
| Memory-mapped FASTA | – | – | Yes (.fai indexed) |
| Parallelism | Fork-based | Lambda | Thread-based (rayon) |
| Web interface | REST API | – | Built-in GUI + REST API |
| Installation | Complex | dotnet | cargo build --release |
| Binary size | ~200 MB | ~50 MB | 3.3 MB |
| Dependencies | >10 CPAN modules | .NET runtime | None |
| VEP field accuracy | Reference | – | 100% (23/23 fields) |

## 4. Discussion

### 4.1 Performance Implications

On the GIAB HG002 gold-standard dataset, fastVEP annotates a complete human WGS 22-24x faster than single-threaded Ensembl VEP (Table 3), and finishes the same genome in about 93 seconds, roughly 44,000 variants per second, with its default multi-threaded pipeline. The fixed 2.6-second binary-cache startup matters only at small variant counts. As input grows, single-threaded throughput rises to 20,466 variants per second on the full 4.05M-variant WGS, showing that the startup cost amortizes efficiently.

The practical consequences are large. Annotating 500,000 variants takes roughly 24 seconds with single-threaded fastVEP, or about 11 seconds multi-threaded, against approximately 9-10 minutes with single-threaded Ensembl VEP. For population-scale studies processing millions of variants, that is the difference between minutes and hours.

Four optimizations underpin this performance: O(T+E) indexed GFF3 assembly, which loads transcripts 12-80x faster; the binary transcript cache, which eliminates repeated GFF3 parsing; tabix-indexed GFF3 loading, which restricts parsing to variant-overlapping regions; and memory-mapped FASTA, which gives zero-copy reference access. Together they remove the startup-dominated bottleneck that previously limited throughput on real data.

### 4.2 Accuracy Validation

fastVEP matches Ensembl VEP v115.1 on all 23 annotation fields across 2,340 transcript-allele pairs, covering consequence terms, HGVS nomenclature, coordinate positions, amino acid changes, codon changes, splice annotations, gene metadata, and transcript quality indicators (TSL, APPRIS, CANONICAL). Because the validation used real Ensembl GRCh38 gene models and the VEP project’s own example variant set, it is strong evidence of annotation fidelity.

The consequence type distribution is identical between the two tools for all 12 consequence types observed in the validation dataset, including rare categories such as splice_polypyrimidine_tract_variant and NMD_transcript_variant.

### 4.3 Extended Splice Site Detection

fastVEP reports six categories of splice site variant, against the three Ensembl VEP traditionally reports. The splice_donor_5th_base_variant, splice_donor_region_variant, and splice_polypyrimidine_tract_variant terms are defined in the Sequence Ontology but often omitted by annotation tools, and they classify variants near splice sites more precisely. This matters most for interpreting variants of uncertain significance in clinical genomics, where splice disruption is a common mechanism of pathogenicity (Jaganathan et al., 2019).

### 4.4 Architectural Design Decisions

Four design decisions deserve comment.

**Modular crate structure.** The twelve-crate workspace allows components to be compiled and tested independently, which speeds development and improves code quality. The trait-based provider architecture (TranscriptProvider, SequenceProvider, VariationProvider, AnnotationProvider, GeneAnnotationProvider) lets data sources be substituted and composed without modifying the prediction engine. The fastvep-sa crate, added for clinical database integration, follows the same pattern: annotation providers are discovered at runtime from .osa files in the --sa-dir directory and plugged into the parallel annotation loop automatically.

**Indexed transcript lookup.** Transcript overlap queries use per-chromosome sorted arrays with binary search, following the chunked binary search strategy of echtvar (Pedersen, 2023). Query complexity is O(log n + k) rather than the O(n) linear scan of naive implementations, which is essential for gene models holding tens of thousands of transcripts per chromosome.

**GFF3 as primary annotation source with binary caching.** Ensembl VEP relies on pre-built caches in Perl’s Storable or Sereal serialization format. fastVEP instead uses standard GFF3 files as its primary transcript source, which removes the dependency on Perl-specific serialization and uses a widely supported format. To avoid the cost of reparsing GFF3 on every run, fastVEP caches transcripts with bincode serialization and gzip compression: the first run parses GFF3 and writes the cache, and later runs load from it in approximately 1.4 seconds for yeast to 2.6 seconds for the full human genome. When a tabix index is available for the GFF3 file, fastVEP loads only transcripts overlapping the input variants, cutting startup time and memory use further.

**47-field CSQ output.** fastVEP writes 47 annotation fields in its CSQ format, against VEP’s 30-field default. The additional fields (CANONICAL, CCDS, ENSP, MANE, MANE_SELECT, MANE_PLUS_CLINICAL, SIFT, PolyPhen, AF, CLIN_SIG, SOMATIC, PHENO, PUBMED, HGVS_OFFSET, and the regulatory motif fields) supply the data most downstream analyses need in a single annotation pass.

**Hybrid web architecture.** The web interface supports both server-side and client-side annotation. Served by the fastVEP binary, requests are processed by the Rust backend with the same performance and accuracy as the CLI, and browser-uploaded GFF3 files are parsed server-side, avoiding the limits of JavaScript parsing on large annotation files. For static deployments such as GitHub Pages, the interface falls back transparently to a client-side JavaScript engine. That fallback allows offline use after page load, at the cost of reduced performance and the maintenance burden of mirroring the Rust logic.

## 5. Methods

### 5.1 Implementation

fastVEP was implemented in Rust 1.94.1 using the 2021 edition, as a Cargo workspace with twelve member crates. External dependencies are limited to: clap (CLI parsing), serde/serde_json/bincode (serialization), anyhow/thiserror (error handling), flate2 (gzip decompression), zstd (zstandard block compression for the v1 fastSA format), zip and lru (the v2 .osa2 chunked-archive format and its chunk cache), log/env_logger (logging), memmap2 (memory-mapped I/O), noodles-bgzf/noodles-tabix/noodles-csi/noodles-core (tabix-indexed file access), rayon and crossbeam (parallelism), and tempfile (atomic output). The Bloom filters used by the .osa2 format are implemented in-crate with no external dependency. The optional standalone web server (fastvep-web) additionally uses axum/tokio/tower. No C libraries or foreign function interfaces are used.

### 5.2 Benchmarking

All benchmarks ran on an Apple M-series ARM64 system under macOS Darwin 24.6.0. The fastVEP binary was compiled with --release (full optimizations). Each benchmark was run three times and the median time reported.

Throughput benchmarks used the GIAB HG002 v4.2.1 high-confidence benchmark VCF (4,048,342 variants) annotated against full Ensembl GRCh38 release 115 GFF3 annotations (508,530 transcripts) with the GRCh38 primary-assembly FASTA reference and HGVS nomenclature enabled. fastVEP started from its binary transcript cache (bincode plus gzip).

For the head-to-head comparison, both tools ran single-threaded on an Apple M4, with fastVEP set to RAYON_NUM_THREADS=1. Ensembl VEP v115.1 (Docker image ensemblorg/ensembl-vep:release_115.1) ran in --gff mode against a bgzip-compressed, tabix-indexed copy of the same GRCh38 release 115 GFF3, with matching options (--vcf --hgvs --symbol --canonical --no_stats). In the supplementary-annotation conditions, ClinVar and gnomAD were supplied to fastVEP as native fastSA (.osa/.osa2) databases and to Ensembl VEP as --custom tabix-indexed VCFs built from identical source data.

A single-process whole-genome VEP run exhausts the default Docker memory of 7.65 GiB above approximately 2.7 million variants. VEP’s genome-wide time was therefore obtained by running each chromosome (1-22) separately and summing, with every per-chromosome time taken as the minimum of two independent runs to remove transient system contention. The chromosome 22 scaling series (1,000 / 5,000 / 10,000 / 50,284 variants) and all fastVEP timings are medians of three runs. Ensembl VEP rejects an unindexed GFF3 outright in --gff mode, so no unindexed-GFF configuration was benchmarked.

Supplementary-annotation format benchmarks built the v1 (.osa) and v2 (.osa2) representations of ClinVar (whole database) and REVEL (chromosome 1) from their standard source files with fastvep sa-build, recording on-disk size and build time. Annotation output was compared by md5 over the annotated VCF of a 673,000-variant real call set.

Cross-organism benchmarks used full Ensembl release 115 GFF3 annotations and FASTA references for yeast (*S. cerevisiae* R64-1-1, 7,036 transcripts), Drosophila (BDGP6.54, 35,442 transcripts), Arabidopsis (TAIR10, 54,013 transcripts), mouse (GRCm39, 142,626 transcripts), and human (GRCh38, 508,530 transcripts). Gold-standard variant sources were Ensembl/SGD variation (yeast), DGRP2 (Drosophila), 1001 Genomes (Arabidopsis), Ensembl/EVA variation (mouse), and GIAB HG002 v4.2.1 (human).

Binary size was measured on the release-compiled, unstripped executable.

### 5.3 Accuracy Validation

Field-level accuracy was validated by running both fastVEP and Ensembl VEP v115.1 (Docker image ensemblorg/ensembl-vep:release_115.1) on the same 173-variant VEP example dataset with equivalent parameters (--gff, --fasta, --hgvs, --symbol, --canonical). The compare_vep.py script parsed CSQ fields from both VCF outputs, matched annotations by (allele, transcript) pair, and compared 23 annotation fields. All fields matched on all 2,340 shared pairs.

### 5.4 Testing

Tests were written with Rust’s built-in test framework (#[test] annotations). Integration tests for VEP compatibility were placed in a separate file (vep_compat.rs) and validated against patterns from Ensembl VEP’s Parser_VCF.t test suite and the test VCF files distributed with VEP release 115. The 47-field CSQ output format was validated against expected field values for missense, frameshift, downstream, and intronic variants. The standard genetic code implementation was validated by testing all 64 codons against NCBI translation table 1.

### 5.5 Availability

fastVEP is open source under the Apache License 2.0. Source code, documentation, and pre-built test data are available at https://github.com/Huang-lab/fastVEP. The tool builds from source with cargo build --release on any platform with a Rust toolchain (Linux, macOS, Windows). A hosted web server for interactive variant annotation runs at https://fastVEP.org; as of July 2026 it had served 2,372 genome analysis sessions and annotated 19,013 variants.

