## Supplementary Information for "fastVEP: A High-Performance Variant Effect Predictor Implemented in Rust"

### Supplementary Materials

#### Supplementary Table S1. Complete list of 49 SO consequence terms implemented in fastVEP.

| **Rank** | **SO Term** | **Impact** | **Description** |
| --- | --- | --- | --- |
| 1 | transcript_ablation | **HIGH** | Complete loss of transcript |
| 2 | splice_acceptor_variant | **HIGH** | Last 2 bases of intron (3’ end) |
| 3 | splice_donor_variant | **HIGH** | First 2 bases of intron (5’ end) |
| 4 | stop_gained | **HIGH** | Premature stop codon |
| 5 | frameshift_variant | **HIGH** | Indel not divisible by 3 in CDS |
| 6 | stop_lost | **HIGH** | Loss of stop codon |
| 7 | start_lost | **HIGH** | Loss of start codon (ATG) |
| 8 | transcript_amplification | **HIGH** | Transcript amplification |
| 9 | feature_elongation | **MODIFIER** | Feature elongation |
| 10 | feature_truncation | **MODIFIER** | Feature truncation |
| 11 | inframe_insertion | **MODERATE** | Insertion divisible by 3 in CDS |
| 12 | inframe_deletion | **MODERATE** | Deletion divisible by 3 in CDS |
| 13 | missense_variant | **MODERATE** | Different amino acid |
| 14 | protein_altering_variant | **MODERATE** | Protein sequence change (complex) |
| 15 | splice_region_variant | **LOW** | 3–8 bp into intron or 1–3 bp into exon |
| 16 | splice_donor_5th_base_variant | **LOW** | 5th intronic base from donor |
| 17 | splice_donor_region_variant | **LOW** | Positions 3–6 from donor |
| 18 | splice_polypyrimidine_tract_variant | **LOW** | 3–15 bases from acceptor |
| 19 | incomplete_terminal_codon_variant | **LOW** | Incomplete codon at CDS end |
| 20 | start_retained_variant | **LOW** | Start codon preserved |
| 21 | stop_retained_variant | **LOW** | Stop codon preserved |
| 22 | synonymous_variant | **LOW** | Same amino acid |
| 23 | coding_sequence_variant | **MODIFIER** | CDS catch-all |
| 24 | mature_miRNA_variant | **MODIFIER** | In mature miRNA |
| 25 | 5_prime_UTR_variant | **MODIFIER** | In 5’ UTR |
| 26 | 3_prime_UTR_variant | **MODIFIER** | In 3’ UTR |
| 27 | non_coding_transcript_exon_variant | **MODIFIER** | In non-coding exon |
| 28 | intron_variant | **MODIFIER** | In intron |
| 29 | NMD_transcript_variant | **MODIFIER** | In NMD transcript |
| 30 | non_coding_transcript_variant | **MODIFIER** | In non-coding transcript |
| 31 | coding_transcript_variant | **MODIFIER** | In coding transcript |
| 32 | upstream_gene_variant | **MODIFIER** | Within 5 kb upstream |
| 33 | downstream_gene_variant | **MODIFIER** | Within 5 kb downstream |
| 34 | TFBS_ablation | **HIGH** | TF binding site ablation |
| 35 | TFBS_amplification | **MODERATE** | TF binding site amplification |
| 36 | TF_binding_site_variant | **MODIFIER** | In TF binding site |
| 37 | regulatory_region_ablation | **HIGH** | Regulatory region ablation |
| 38 | regulatory_region_amplification | **MODERATE** | Regulatory region amplification |
| 39 | regulatory_region_variant | **MODIFIER** | In regulatory region |
| 40 | intergenic_variant | **MODIFIER** | Between genes |
| 41 | sequence_variant | **MODIFIER** | Generic sequence variant |
| 42 | copy_number_change | **MODIFIER** | Copy number variation (unspecified) |
| 43 | copy_number_increase | **MODIFIER** | Copy number gain |
| 44 | copy_number_decrease | **MODIFIER** | Copy number loss |
| 45 | short_tandem_repeat_change | **MODIFIER** | STR length change |
| 46 | short_tandem_repeat_expansion | **MODIFIER** | STR expansion |
| 47 | short_tandem_repeat_contraction | **MODIFIER** | STR contraction |
| 48 | unidirectional_gene_fusion | **MODIFIER** | Gene fusion from breakend |
| 49 | transcript_variant | **MODIFIER** | Generic SV overlap with transcript |
